# Comparative structural dynamic analysis of GTPases

**DOI:** 10.1101/370197

**Authors:** Hongyang Li, Xin-Qiu Yao, Barry J. Grant

## Abstract

GTPases regulate a multitude of essential cellular processes ranging from movement and division to differentiation and neuronal activity. These ubiquitous enzymes operate by hydrolyzing GTP to GDP with associated conformational changes that modulate affinity for family-specific binding partners. There are three major GTPase superfamilies: Ras-like GTPases, heterotrimeric G proteins and protein-synthesizing GTPases. Although they contain similar nucleotide-binding sites, the detailed mechanisms by which these structurally and functionally diverse superfamilies operate remain unclear. Here we compare and contrast the structural dynamic mechanisms of each superfamily using extensive molecular dynamics (MD) simulations and subsequent network analysis approaches. In particular, dissection of the cross-correlations of atomic displacements in both the GTP and GDP-bound states of Ras, transducin and elongation factor EF-Tu reveals analogous dynamic features. This includes similar dynamic communities and subdomain structures (termed lobes). For all three proteins the GTP-bound state has stronger couplings between equivalent lobes. Network analysis further identifies common and family-specific residues mediating the state-specific coupling of distal functional sites. Mutational simulations demonstrate how disrupting these couplings leads to distal dynamic effects at the nucleotide-binding site of each family. Collectively our studies extend current understanding of GTPase allosteric mechanisms and highlight previously unappreciated similarities across functionally diverse families.

**Author Summary:** GTPases are a large superfamily of essential enzymes that regulate a variety of cellular processes. They share a common core structure supporting nucleotide binding and hydrolysis, and are potentially descended from the same ancestor. Yet their biological functions diverge dramatically, ranging from cell division and movement to signal transduction and translation. It has been shown that conformational changes through binding to different substrates underlie the regulation of their activities. Here we investigate the conformational dynamics of three typical GTPases by *in silico* simulation. We find that these three GTPases possess overall similar substrate-associated dynamic features, beyond their distinct functions. Further identification of key common and family-specific elements in these three families helps us understand how enzymes are adapted to acquire distinct functions from a common core structure. Our results provide unprecedented insights into the functional mechanism of GTPases in general, which potentially facilitates novel protein design in the future.

## Introduction

Guanosine Triphosphate Phosphohydrolases (GTPases) are ubiquitous molecular machines mediating a variety of essential cellular processes [1]. Harnessing the GTP hydrolysis to modulate the affinity of partner molecule binding, GTPases transduce intracellular signals, control cell division and differentiation, and direct protein synthesis and translocation [2–5]. In general, GTP-bound GTPases in the active state are able to interact with partner effectors and regulate effector-mediated processes. GTP hydrolysis leads to the dissociation of GTPases from effectors, whereas exchange of GDP for GTP activates GTPases and restarts the signaling or protein synthesis cycle [6,7]. Two classes of accessory proteins are involved in regulating this reaction cycle. GTPase-activating proteins (GAPs) accelerate the GTPase activity and the inactivation of GTPases, whereas guanine nucleotide exchange factors (GEFs) promote GDP dissociation and subsequent GTP binding, activating GTPases [8–10].

There are three major GTPase superfamilies: monomeric small Ras-like GTPase, heterotrimeric G protein α subunit (Gα) and protein-synthesizing GTPase. Both small and heterotrimeric G proteins participate in signal transduction. As the primary coupling molecule to membrane receptors, Gα together with its partner βγ subunits (Gβγ) mediate the very early stage signal transduction initiated by extracellular stimuli. In contrast, small GTPase does not interact with receptors directly and regulates more downstream events in the cascade. Finally, the protein-synthesizing proteins participate in initiation, elongation and termination of mRNA translation. Underlying this functional difference are the low sequence identity (<20%) and overall different molecular shapes among these three types of GTPases. In particular, whereas small G protein consists of a single canonical Ras-like catalytic domain (RasD), Gα has an extra α-helical domain (HD) inserted and elongation factor EF-Tu has two extra β-barrel domains (D2 and D3) subsequent to the C-terminus (**Figure 1**). In addition, unlike small G protein that always acts as a monomer, Gα can form a complex with Gβγ and undergoes a cycle of altered oligomeric states during function.

**Figure 1.**
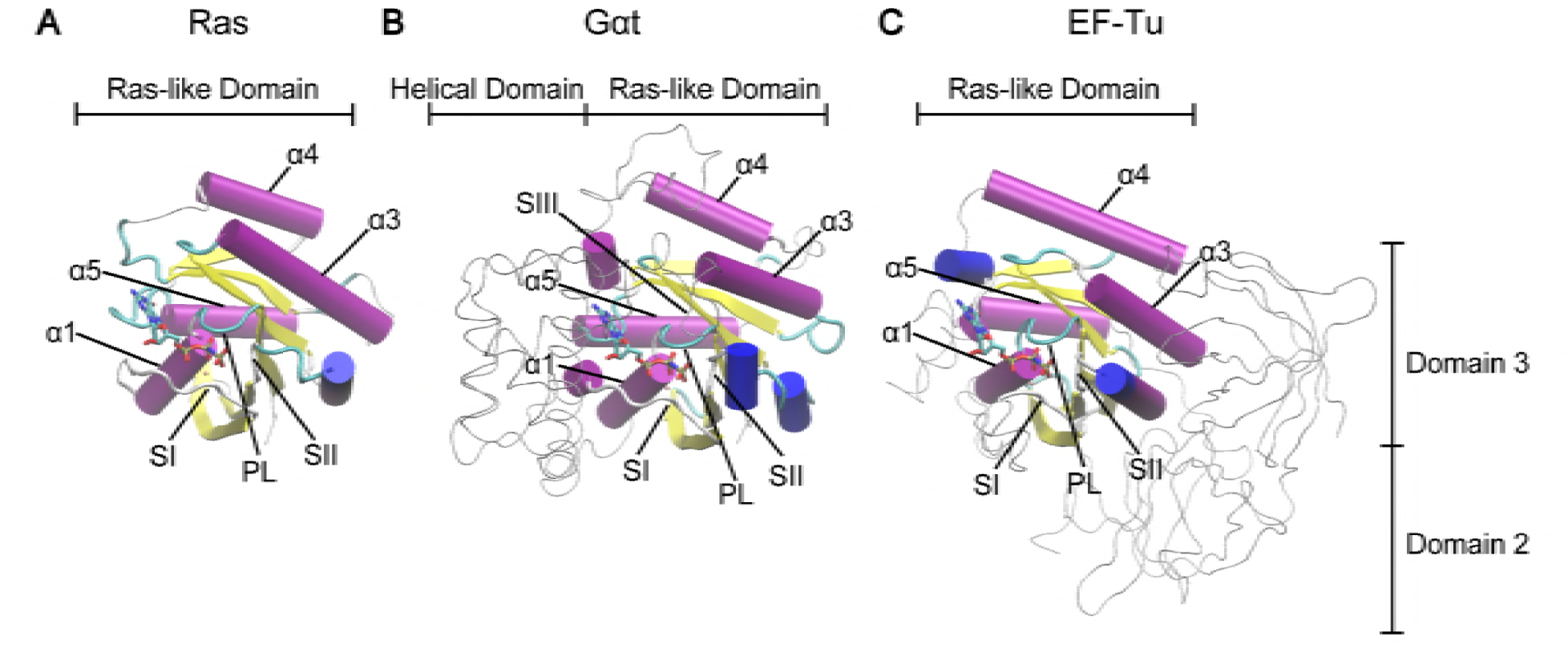
Structural comparison of Ras, Gαt and EF-Tu reveals common canonical Ras-like domain. The Ras-like domains of Ras (**A**), Gαt (**B**) and EF-Tu (**C**) are shown in cartoon and the extra domains in Gαt and EF-Tu are shown as gray tubes. Highly conserved regions (PL, SI, and SII) and helices (α1, α3, α4, and α5) are labeled. The PDB IDs of these three structures are 5P21 (Ras), 1TND (Gαt) and 1TTT (EF-Tu).

In contrast to the functional and structural diversity, GTPases display significant conservation in the core structure of the catalytic domain. Small GTPase, Gα and EF-Tu contain a RasD consisting of six β strands (β1-β6) and five α helices (α1-α5) flanking on both sides of the β sheet (**Figure 1**). Three highly conserved loops named P-loop (PL), switch I (SI), and switch II (SII) constitute the primary sites coordinating the nucleotide phosphates. This structural similarity suggests that at a fundamental level small GTPase, Gα and EF-Tu may utilize the same mode of structural dynamics for their allosteric regulation, which is likely inherited from their common evolutionary ancestor [11,12]. However, it is currently unclear what are the general atomistic mechanisms underlying GTPase allostery and how these common mechanisms can be adapted to have specific function.

Recent computational and experimental studies have gained much insight into the allosteric mechanisms of individual small and heterotrimeric G protein systems. Principal component analysis (PCA) of crystallographic structures and molecular dynamics (MD) simulations characterized the structural dynamics of small GTPase Ras and revealed an intriguing dynamical partitioning of Ras structure into two lobes: the N-terminal nucleotide binding lobe (lobe1) and the C-terminal membrane anchoring lobe (lobe2) [13,14]. Several allosteric sites were identified in lobe 2 or between lobes, including L3 (the loop between β2 and β3), L7 (the loop between α3 and β5), and α5. Importantly, α5 is the major membrane-binding site and has been related to the nucleotide modulated Ras/membrane association [15]. In addition, binding of small molecules at L7 has been reported to affect the ordering of SI and SII [16]. Intriguingly, recent studies of Gα have revealed nucleotide associated conformational change and bilobal substructures in the catalytic domain largely resembling those in Ras [17,18]. The allosteric role of lobe 2, which contains the major binding interface to receptors, has also been well established for Gα [18–27]. Furthermore, the comparison between G proteins and translational factors via sequence and structural analysis indicates a conserved molecular mechanism of GTP hydrolysis and nucleotide exchange, and cognate mutations of key residues in the nucleotide-binding regions showed similar functional effects among these systems [2,6,7,12]. Collectively, these consistent findings from separate studies support the common allosteric mechanism hypothesis of GTPases and underscore a currently missing detailed residue-wise comparison of the structural dynamics among different GTPase superfamilies.

In this study, we compare and contrast the nucleotide-associated conformational dynamics between H-Ras (H isoform of Ras), Gαt (transducin α subunit) and EF-Tu (elongation factor thermo unstable), and describe how this dynamics can be altered by single point mutations in both common and family-specific ways. This entails the application of an updated PCA of crystallographic structures, multiple long time (80-ns) MD simulations, and recently developed network analysis approach of residue cross-correlations [18]. In particular, we identify highly conserved nucleotide dependent correlation patterns across GTPase families: the active GTP-bound state displays stronger correlations both within lobe1 and between lobes, exhibiting an overall “dynamical tightening” consistent with the previous study in Gα alone [18]. Detailed inspection of the residue level correlation networks along with mutational MD simulations reveal several common key residues that are potentially important for mediating the inter-lobe communications. Point mutations of these residues substantially disrupt the couplings around the nucleotide binding regions in Ras, Gαt and EF-Tu. In addition, with the same network comparison analysis, we identify Gαt and EF-Tu specific key residues. Mutations of these residues significantly disrupt the couplings in Gαt and EF-Tu but have no or little effect in Ras. Our results are largely consistent with findings from experimental mutagenesis, with a number of dynamical disrupting mutants have been shown to have altered activities in either Ras or Gα. Our new predictions can be promising targets for future experimental testing.

## Results

### Principal component analysis (PCA) of Ras, Gαt/i and EF-Tu crystallographic structures reveals functionally distinct conformations

Previous PCA of 41 Ras crystallographic structures revealed distinct GDP, GTP and intermediate mutant conformations [13]. Updating this analysis to include the 121 currently available crystallographic structures (**Table S1**) reveals consistent results but with two additional conformations now evident (**Figure 2A**). In addition to GDP (green in **Figure 2A**), GTP (red), and mutant forms, GEF-bound nucleotide free (purple) and so-called ‘state 1’ forms (orange) are now also apparent. In the GEF-bound form, the SI region is displaced in a distinct manner – 12Å away from the nucleotide-binding site coincident with the insertion of a helix of GEF into the PL-SI cleft. The state 1 GTP-bound form was first observed via NMR and later high-resolution crystal structures were solved [28–30]. In contrast to the canonical GTP-bound conformation (red), the state 1 form (orange) lacks interaction between the two switches and the γ-phosphate of GTP, resulting in a moderate 7Å displacement of SI away from its more closed GTP conformation.

**Figure 2.**
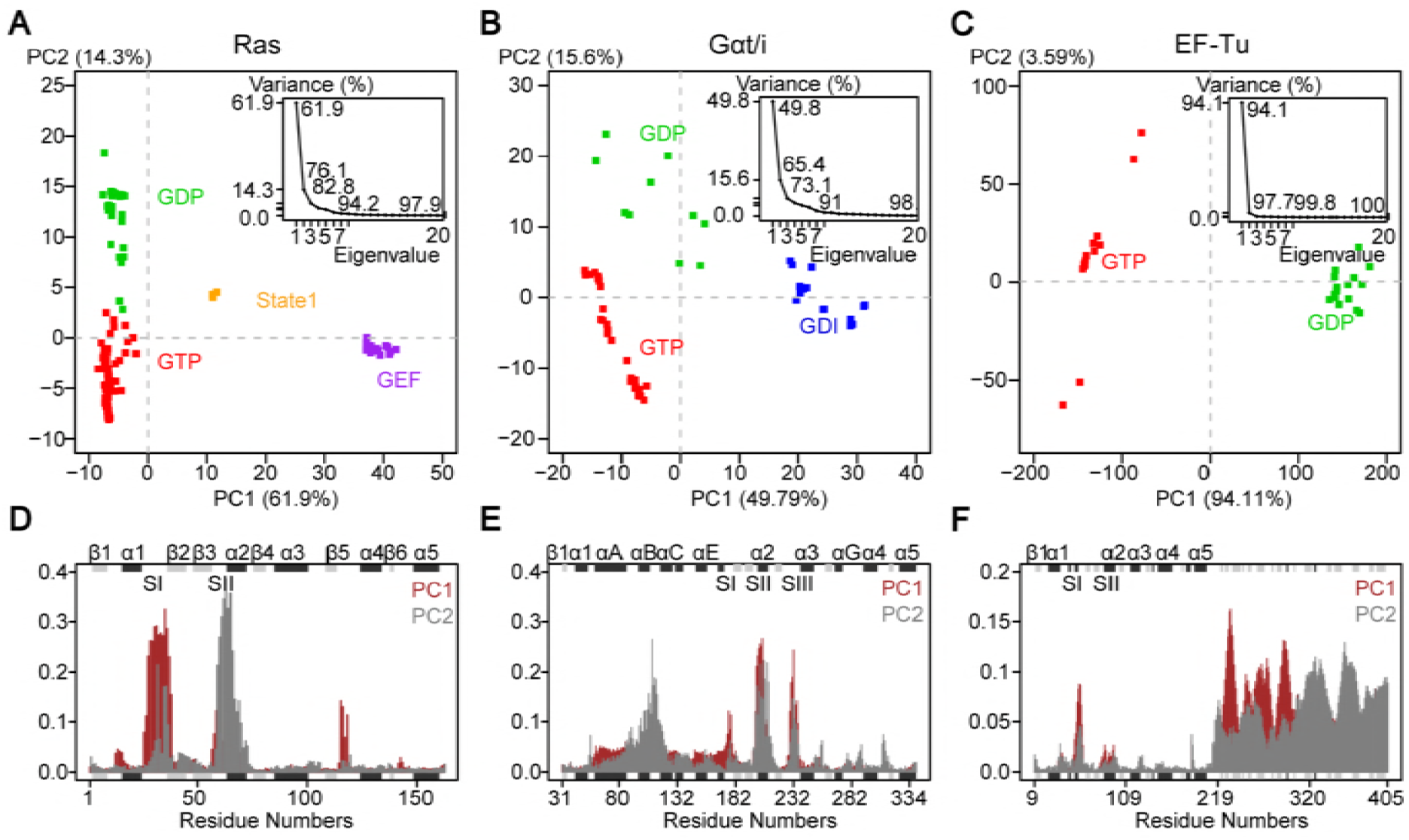
Principal component analysis of Ras, Gαt/i and EF-Tu crystallographic structures reveals distinct nucleotide-associated conformations. (**A-C**) Projection of 121 Ras (**A**), 53 Gαt/i (**B**) and 23 EF-Tu (**C**) PDB structures (represented as squares; see also **Table S1-S3**) onto the first two PCs reveals different conformational clusters corresponding to GTP (red), GDP (green), GEF (purple) and GDI (blue) bound states. A distinct cluster of GTP-bound structures in Ras corresponds to the “State 1” state (orange). The inserted figures show that the first two PCs capture 76.1%, 65.4% and 97.7% of the total structural variances in Ras, Gαt/i and EF-Tu, respectively. (**D-F**) The contributions of each residue to PC1 (brown) and PC2 (grey) show that the switch regions mainly correspond to the accumulated structural differences in Ras (**D**) and Gαt/i (**E**). In addition to switch regions, Domain 2 and Domain 3 also contribute to the structure differences in EF-Tu (**F**). The marginal black and grey rectangles with labels on top of them represent the location of alpha-helix and beta-strand secondary structures.

The first two PCs capture more than 75% of the total mean-square displacement of all 121 Ras structures. Residue contributions from SI and SII dominate PC1 and PC2 (**Figure 2D**). The height of each bar in **Figure 2D** displays the relative contribution of each residue to a given PC. PC1 mainly describes the opening and closing of SI – more open in GEF-bound and state 1 forms, and more closed in nucleotide bound structures. PC1 also captures smaller scale displacement of L8 (the loop between β5 and α4), which resides 5Å closer to the nucleotide-binding pocket in the GEF-bound structures than the GTP-bound structure set. PC2 depicts SII displacements and clearly separates GTP from GDP bound forms (red and green, respectively). As we expect, the lack of γ-phosphate in the GDP releases SII from the nucleotide, whereas in the GTP form SII is fixed by the hydrogen bond of the backbone amide of G60 with the γ-phosphate oxygen atom. This is also shown in the state 1 form where the hydrogen bond is disrupted with SII moderately displaced from the nucleotide (4Å on average from the canonical GTP group structures).

PCA of 53 available Gαt/i structures described recently (**Table S2**) revealed three major conformational groups: GTP (red in **Figure 2B**), GDP (green) and GDI (GDP dissociation inhibitor; blue) bound forms [18]. The first two PCs capture over 65% of the total variance of Cα atom positions in all structures. The dominant motions along PC1 and PC2 are the concerted displacements of SI, SII and SIII in the nucleotide-binding region as well as a relatively small-scale rotation of the helical domain with respect to RasD (**Figure 2E**).

PC1 separates GDI-bound from non-GDI bound forms. In GDI-bound structures the GDI interacts with both the HD and the cleft between SII and SIII of the Ras-like domain, increasing the distance between SII and SIII. Similar to Ras, PC2 of Gαt/i clearly distinguishes the GTP and GDP-bound forms, where again the unique γ-phosphate (or equivalent atom in GTP analogs) coordinates SI and SII. In addition, the SIII is displaced closer to the nucleotide, effectively closing the nucleotide-binding pocket.

PCA of 23 available full-length EF-Tu structures reveals distinct GTP and GDP conformations (**Table S3**). PC1 dominantly captures nearly 95% of the total structural variance of Cα atom positions (**Figure 2C**). It mainly describes the dramatic conformational transition in SI as well as the large rotation of two β-barrel domains D2 and D3 (**Figure 2F**). In the GTP-bound form, the C-terminal SI is coordinated to the γ-phosphate and, Mg^2+^ ion, forming a small helix near SII. Meanwhile, D2 and D3 are close to RasD and create a narrow cleft with SI, serving as the binding site for tRNA [31]. In the GDP-bound form, the C-terminal helix in SI unwinds and forms a β-hairpin, protruding towards D2 and D3 [32]. The highly conserved residue T62 (T35 in Ras) of EF-Tu moves more than 10Å away from its position in the GTP form and loses interaction with the Mg^2+^ ion. In addition, D3 rotates towards SI and D2 moves far away from the Ras-like domain. In contrast to PC1, PC2 only captures a very small portion (3.59%) of the structural variance in EF-Tu (**Figure 2F**). The major conformational change along PC2 is a small-scale rotation of D2 and D3 with respect to RasD in the GTP form.

PCA of Ras, Gαt/i and EF-Tu demonstrates that the binding of different nucleotides and protein partners can lead to a rearrangement of global conformations in a consistent manner. In particular, within RasD, these three families display conserved nucleotide-dependent conformational distributions with major contributions from the switch regions. In the GTP-bound form of these proteins, SI and SII are associated with the nucleotide through interacting with γ-phosphate. Despite these similarities, critical questions about their functional dynamics remain unanswered: How does nucleotide turnover lead to allosteric regulation of distinct partner protein-binding events? To what extent are the structural dynamics of these proteins similar beyond the switch region displacements evident in accumulated crystal structures? How do distal disease-associated mutations affect the functional dynamics for each family and are there commonalities across families? In the next section, we report MD simulations that address these questions, which are not answered by accumulated static experimental structures.

### MD simulations reveal distinct nucleotide-associated flexibility and cross-correlation near functional regions

MD simulations reveal distinct nucleotide-associated flexibility at known functional regions. Representatives of the distinct GTP and GDP-bound conformations of Ras, Gαt and EF-Tu were selected as starting points for MD simulation. Five replicated 80-ns MD simulations of these three proteins for each state (GTP and GDP totaling 2.4us; see **Materials and Methods**) exhibit high flexibility in the SI, SII, SIII/α3 and loop L3, L7, L8 and L9 regions (**Figure 3A-C**). The Cα atom root-mean-square fluctuation (RMSF) in Gαt shows that SI is significantly more flexible in the GDP-bound state (**Figure 3B**). The C-terminal SI of Ras and EF-Tu, corresponding to the shorter SI in Gαt, is also more flexible with GDP bound (**Figure 3A & C**). Interestingly, the middle part of SI in Ras and EF-Tu show higher fluctuations in the GTP-bound state. Moreover, SII is more flexible in the GTP-bound state in Ras. Detailed inspection reveals that SII always stays away from the nucleotide during the GDP-bound state MD simulations, whereas SII sometimes moves close to and interacts with the unique γ-phosphate of GTP, leading to higher flexibility in the GTP-bound state. In contrast, the flexibility of SII in Gαt has no significant difference between states, whereas SII in EF-Tu is less flexible with GTP bound. This is due to the relatively compact interactions between SII and the unique D2 and D3 in the GTP-bound EF-Tu. In fact, D2 and D3 show extremely higher flexibility in the GDP state (**Figure 2C**). Overall, the nucleotide-dependent flexibility of RasD in Ras, Gαt and EF-Tu are quite similar except for SII.

**Figure 3.**
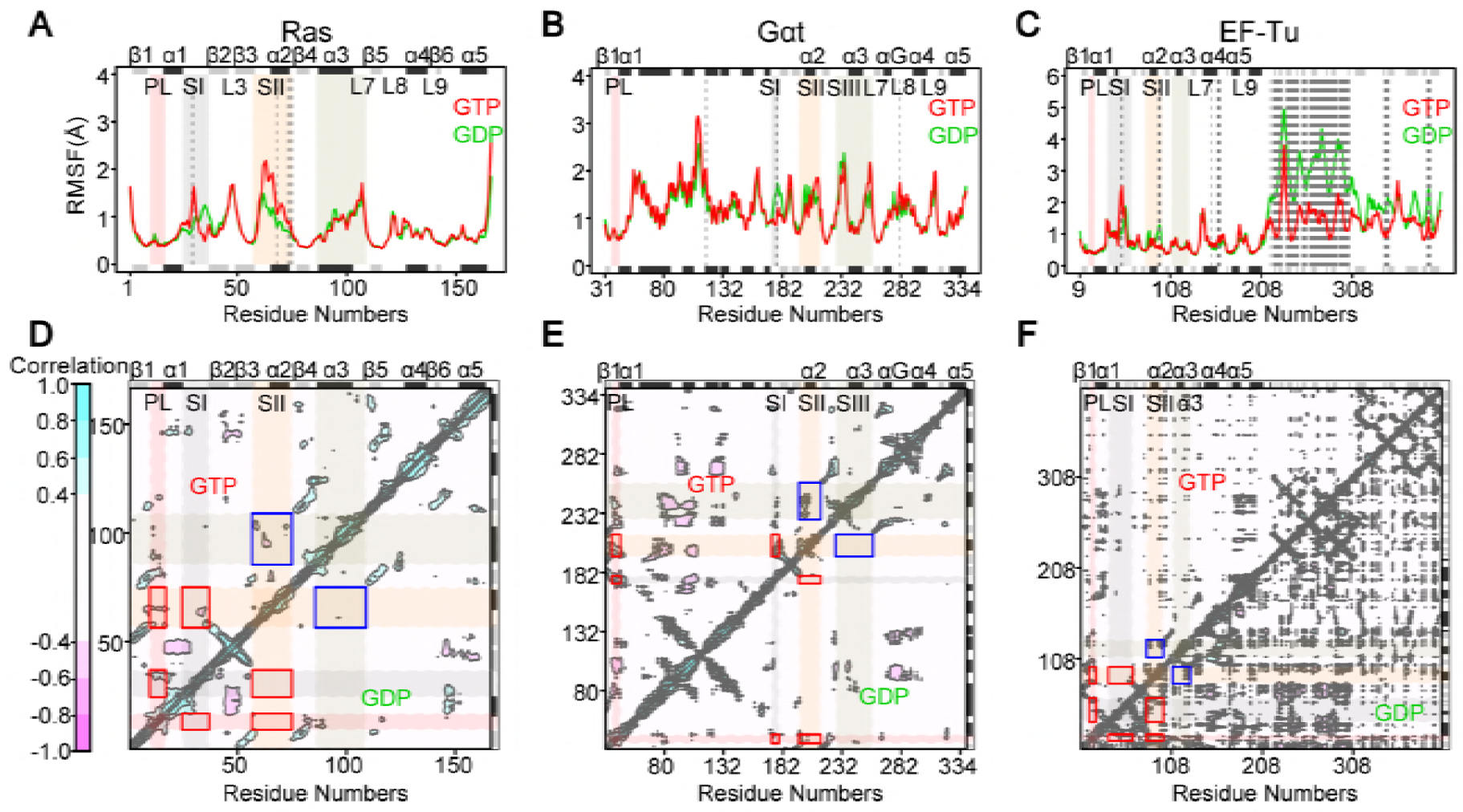
Nucleotide specific residue fluctuations and cross-correlations of atomic displacements from molecular dynamics simulations. (**A-C**) The ensemble averaged root-mean-square fluctuation (RMSF) reveals nucleotide dependent flexibilities that are consistent in the Ras-like domain of Ras (**A**), Gαt (**B**) and EF-Tu (**C**). Residues with significant differences (*p*-value < 0.01) between GTP and GDP bound states are highlighted with dashed lines. (**D-F**) The cross-correlations reveal stronger intra-lobe1 couplings between PL, SI and SII (red rectangles) and inter-lobe couplings between SII and SIII/α3 (blue rectangles) in the GTP-bound state (upper triangle) for both Ras (**D**) and Gαt (**E**).

The cross-correlations of atomic displacements derived from MD simulations also manifest conserved nucleotide-associated coupling in these three systems (**Figure 3D-F**). In both Ras and Gαt, significantly stronger couplings within the catalytic lobe 1 between PL, SI and SII can be found only in the GTP-bound state (red rectangles in **Figure 3D & E**). Interestingly, a unique inter-lobe coupling between SII and SIII/α3 also characterizes the GTP-bound state in both systems (blue rectangles in **Figure 3D & E**). In EF-Tu, the intra-lobe 1 and inter-lobe couplings are similar between states (red and blue rectangles in **Figure 3F**). Intriguingly, a lot of negative correlations between D2 and RasD of EF-Tu are found in the GDP-bound state, indicating the swing motion of D2 with respect to RasD during MD simulations (lower triangle in **Figure 3F**).

### Correlation network analysis displays similar nucleotide-associated correlation in Ras, Gαt and ET-Tu

Consensus correlation networks for each nucleotide state were constructed from the corresponding replicate MD simulations. In these initial networks, each node is a residue linked by edges whose weights represent their respective correlation values averaged across simulations (see **Materials and Methods**). These residue level correlation networks underwent hierarchical clustering to identify groups of residues (termed communities) that are highly coupled to each other but loosely coupled to other residue groups. Nine communities were identified for Ras and eleven for Gαt and EF-Tu (**Figure 4**). The two additional family specific communities not present in Ras correspond to two regions of HD in Gαt and D2 and D3 in EF-Tu.

**Figure 4.**
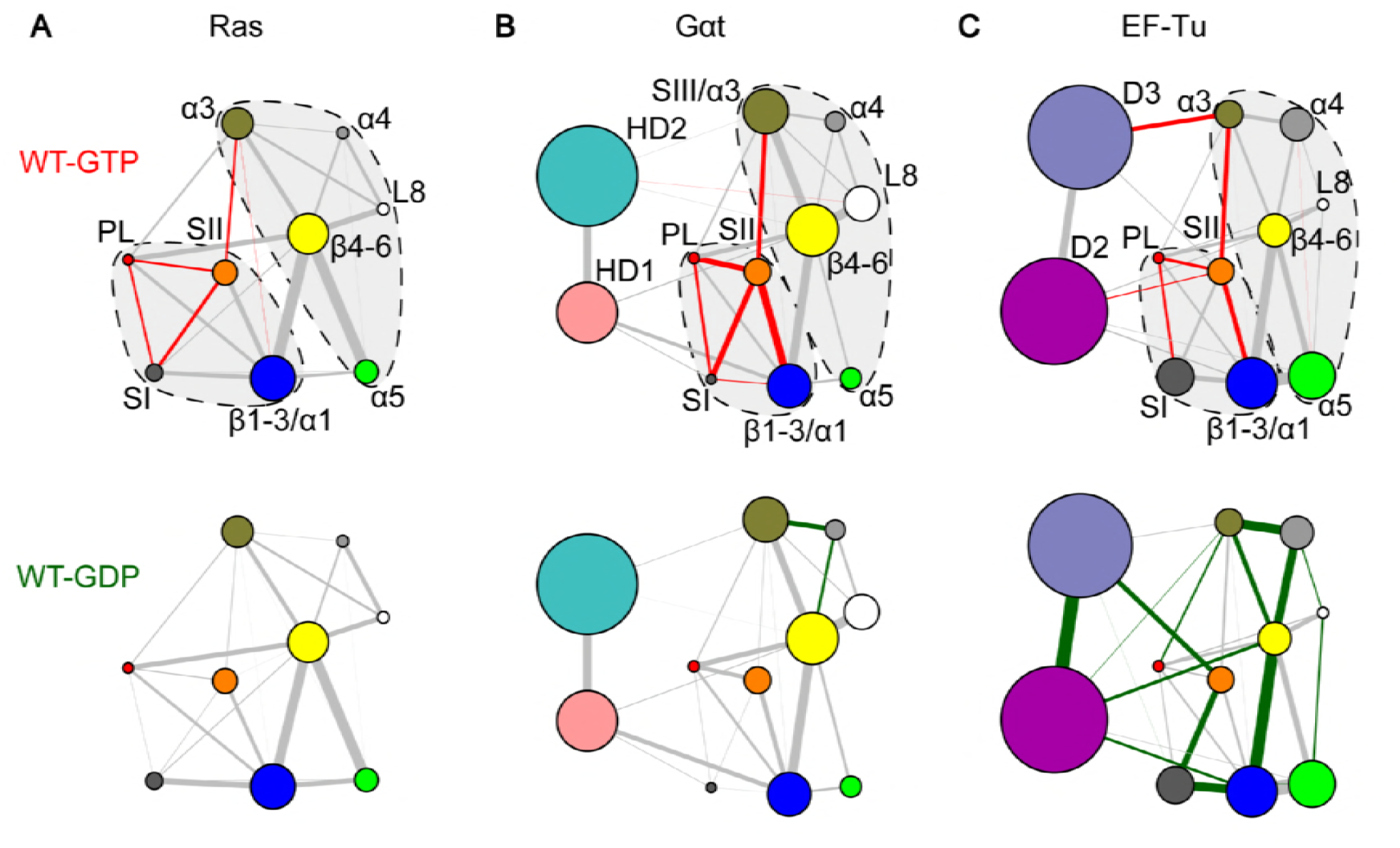
Correlation network analysis reveals similar patterns of nucleotide-dependent couplings in Ras, Gαt and EF-Tu. (**A**) Network communities are represented as colored circles with different radius indicating the number of residues within the community. The width of an edge is determined by the summation of all residue level correlation values between two connected communities. Red and green edges indicate enhanced GTP or GDP couplings that are significantly (*p*-value < 0.05) or more than two-fold stronger in one state than the other. All other lines are colored gray. Dashed lines with a light gray background represent the two-lobe substructures. (**B & C**) Similar nucleotide-associated network patterns are evident in the GTP (top) and GDP (bottom) bound state of Gαt (**B**) and EF-Tu (**C**), except for the SI and SII coupling.

In the resulting community networks the width of an edge connecting two communities is the sum of all the underlying residue correlation values between them. Interestingly, Ras, Gαt and EF-Tu community networks can be partitioned into two major groups (dashed lines in **Figure 4**) corresponding to the previously identified lobes for Ras and the RasD in Gαt [13,18]. The boundary between lobes is located at the loop between α2 and β4. In these proteins, lobe1 includes the nucleotide-binding communities (PL, SI and SII) as well as the N-terminal β1-β3 and α1 structural elements. Lobe2 includes α3-α5, L8 and the C-terminal β4-β6 strands.

Comparing the GTP and GDP community networks of these three proteins reveals common nucleotide-dependent coupling features. In particular, for Ras and Gαt, comparing the relative strength of inter-community couplings in GTP and GDP networks using a nonparametric Wilcoxon test across simulation replicates reveals common significantly distinct coupling patterns (colored edges in **Figure 4A & B**). Within lobe1 stronger couplings between PL, SI and SII are observed for the GTP state of both families. This indicates that the γ-phosphate of GTP leads to enhanced coupling of these proximal regions. This is consistant with our PCA results above, where PC2 clearly depicts the more closed conformation of SI and SII in the GTP bound structures (**Figure 2D & E**). In addition, a significantly stronger inter-lobe correlation between SII and α3 is evident for the GTP state of both families, which is not available from analysis of the static experimental ensemble alone. This indicates that nucleotide turnover can lead to distinct structural dynamics not only at the immediate nucleotide-binding site in lobe 1 but also at the distal lobe 2 region.

Intriguingly, similar patterns of intra and inter-lobe dynamic correlations are observed in EF-Tu (**Figure 4C**). Within lobe1, significantly stronger correlations between PL-SI and PL-SII are evident in the GTP state, although SI-SII coupling becomes weaker in this state. In fact, the C-terminal β-hairpin of SI moves towards and interacts extensively with SII and D3 in the GDP bound state, leaving the nucleotide-binding site widely open. Moreover, our results reveal that SII and SIII/α3 of EF-Tu are more tightly coupled in the GTP state, resembling the strong inter-lobe couplings in the GTP bound Ras and Gαt. It is worth noting that this conserved structural dynamic coupling is evident only from the comparative network analysis and is not accessible from PCA of crystal structures.

### The common residue-wise determinants of structural dynamics in Ras, Gαt and EF-Tu

Comparative network analysis highlights the common residue-wise determinants of nucleotide-dependent structural dynamics. Besides correlations within lobe1, inter-lobe couplings are also significantly stronger in the GTP state networks of Ras, Gαt and EF-Tu. Inspection of the residue-wise correlations between communities reveals common major contributors to the SII – α3 couplings in the three proteins (red residues in **Table S4**). In particular, M72^Ras^ in SII and V103^Ras^ in α3 act as primary contributors to inter-lobe correlations in Ras. Interestingly, the equivalent residues in the other two systems, F211^Gαt^ or I93^EF-Tu^ in SII and F255^Gαt^ or V126^EF-Tu^ in α3/SIII also contribute to the inter-lobe couplings. We further examined the importance of these residues by MD simulations of mutant GTP-bound systems. Results indicate that each single mutation M72A^Ras^ and V103A^Ras^ can significantly reduce the couplings between SI and PL, indicating that these mutations disturb couplings at distal sites of known functional relevance (**Figure 5A & D**). Moreover, the cognate mutations F211A^Gαt^ and F255^Gαt^ in Gαt not only decouple SI and PL but also SI and SII (**Figure 5B & E**). Similarly, the analogous mutation I93A^EF-Tu^ decreases the correlations between PL and SI, whereas V126A^EF-Tu^ decouples PL and SII (**Figure 5C & F**). The simulation results indicate that single alanine mutation of residues contributing to SII-α3 couplings diminishes the couplings of the nucleotide binding regions, and this allosteric effect is common in all the three proteins.

**Figure 5.**
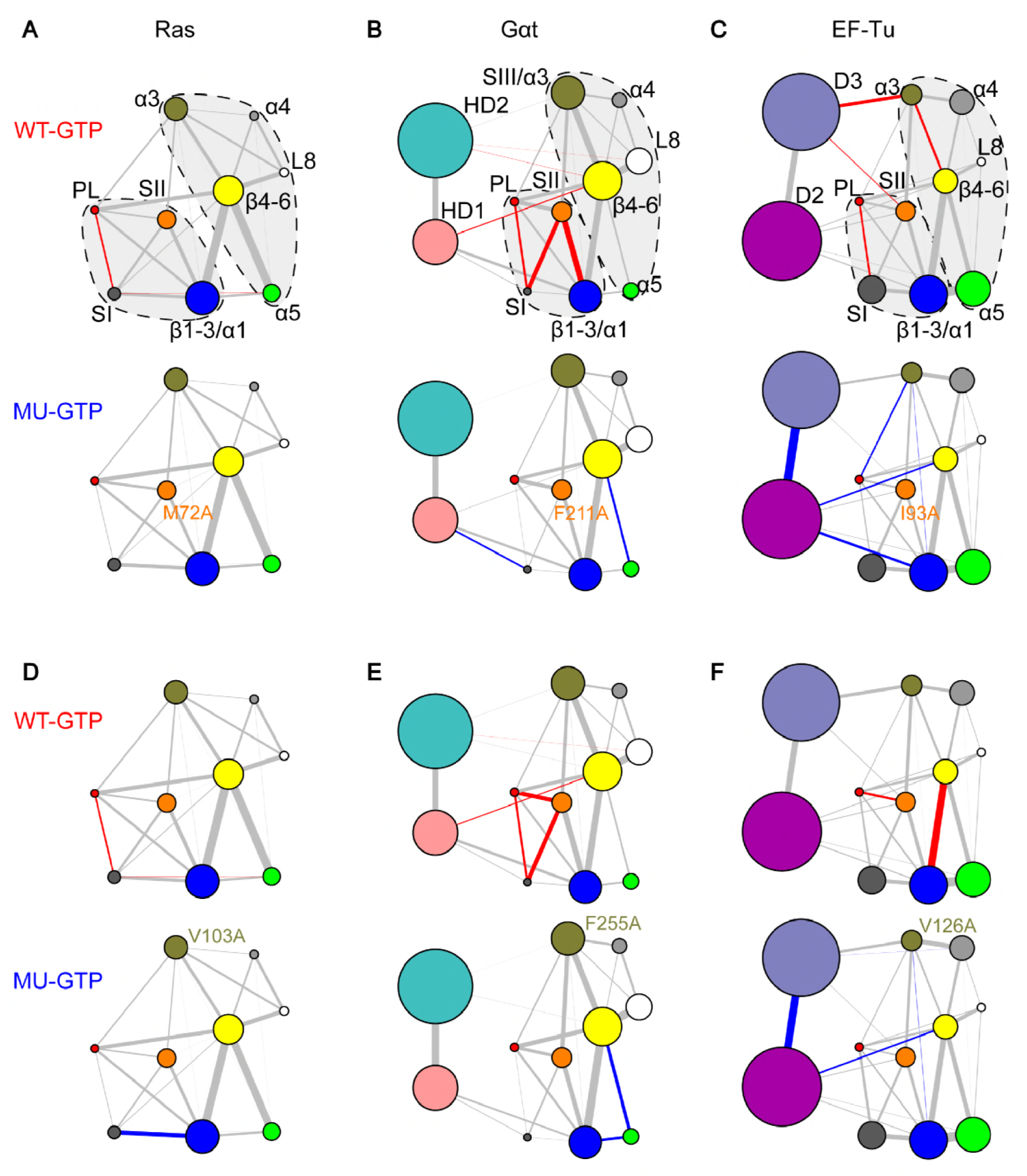
Mutations of common residue-wise determinants of structural dynamics between SII and α3 have similar effects in Ras, Gαt and EF-Tu. Mutations M72A^Ras^ in SII (**A**) and V103A^Ras^ in α3 (**D**) significantly reduce the couplings between PL and SI. The counterpart mutations in Gαt and EF-Tu, F211A^Gαt^ in SII (**B**), F255A^Gαt^ in α3 (**E**), I93A^EF-Tu^ in SII (**C**) and V126A^EF-Tu^ in α3 (**F**) have similar effects in the nucleotide-binding region – significantly reducing the coupling between PL, SI and SII.

Inter-lobe couplings that are distal from the nucleotide binding regions are also shown to be critical for the nucleotide dependent dynamics in Ras, Gαt and EF-Tu. By inspecting the residue level couplings between L3 and α5, we identified common distal inter-lobe couplings in the three proteins. Mutational simulations indicate that the substitutions K188A^Gαt^ and D337A^Gαt^ significantly decouple SI from the PL and SII regions (**Figure 6B & E**). Interestingly, the mutations K188A^Gαt^ and D337A^Gαt^ have been reported to cause a 6-fold and 2-fold increase in nucleotide exchange, respectively, but no direct structural dynamic mechanism was established [19]. We further tested mutations of analogous residues in Ras. We considered both D47^Ras^ and E49^Ras^ as the equivalent residues to K188^Gαt^ (due to the longer L3 region of Ras), and R164^Ras^ as the equivalent residue to D337^Gαt^. Both double mutation D47A/E49A^Ras^ and single mutation R164A^Ras^ significantly reduce the correlations between PL and SI (**Figure 6A & D**). We note that the functional consequences of mutating these residues in Ras has been highlighted in a previous study, in which the salt bridges between D47/E49^Ras^ in L3 and R161/R164^Ras^ in α5 were shown to be involved in the reorientation of Ras with respect to the plasma membrane, and enhanced activation of MAPK pathway [15]. Moreover, substitutions of analogous residues R75A^EF-Tu^ (L3) and D207A^EF-Tu^ (α5) also significantly reduce the couplings between PL and SI (**Figure 6C & F**). Our results indicate that the conserved interactions between L3 and α5 are important for maintaining the close coordination of the distal SI, SII and PL around the nucleotide, and this is common to these three proteins.

**Figure 6.**
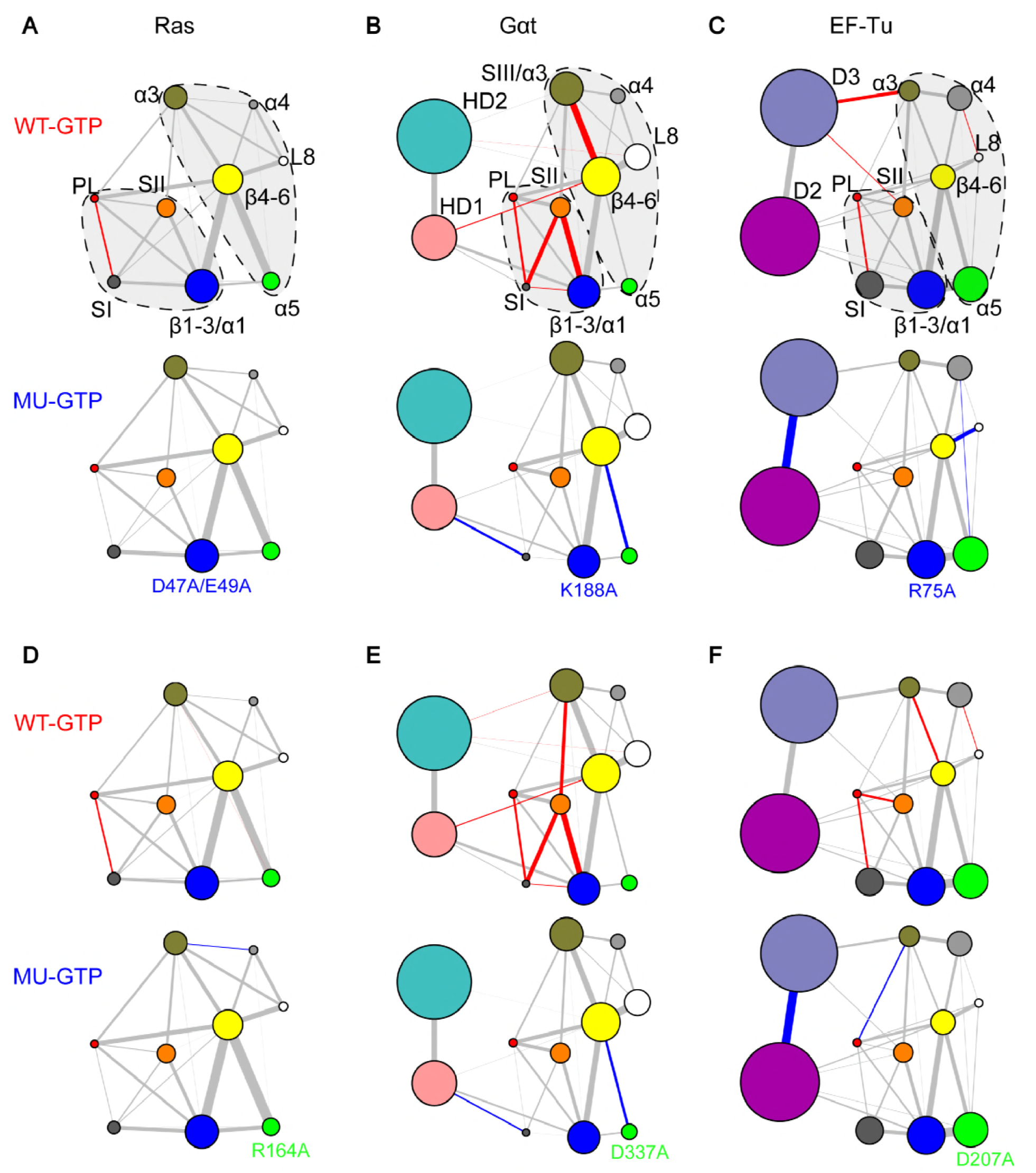
Mutations of common residue-wise determinants of structural dynamics between L3 and α5 have similar effects in Ras, Gαt and EF-Tu. Mutations D47A/E49A^Ras^ in L3 (**A**) and R164A^Ras^ in α5 (**D**) significantly reduce the couplings between PL and SI. The counterpart mutations in Gαt and EF-Tu, K188A^Gαt^ in L3 (**B**), D337A^Gαt^ in α5 (**E**), R75A^EF-Tu^ in L3 (**C**) and D207A^EF-Tu^ in α5 (**F**) have similar effects in the nucleotide-binding region – significantly reducing the coupling between PL, SI and SII.

### Network analysis identifies family-specific residue substitutions that can also perturb structural dynamics

Comparison of the GTP-bound residue-wise networks of Ras, Gαt and EF-Tu reveals that the N-terminus of α3 strongly couples SII only in Gαt and EF-Tu. In particular, we identified residues R201^Gαt^ or A86^EF-Tu^ (SII) and E241^Gαt^ or Q115^EF-Tu^ (α3) as underlying these strong couplings (blue residues in **Table S4**). These residues are specific to Gαt and EF-Tu because the corresponding residues E62^Ras^ in SII and K88^Ras^ in α3 have no contribution in Ras (green residues in **Table S4**). Mutational MD simulations indicate that substitutions E241A^Gαt^ and Q115A^EF-Tu^ have a similar drastic effect on the coupling of nucleotide binding regions (**Figure S1**). In particular, the couplings between PL, SII and PL are all significantly reduced (**Figure S1B & C**). We note that E241A^Gαt^ in Gαs (the α subunit of the stimulatory G protein for adenylyl cyclase) was previously reported to impair GTP binding but the structural basis for this allosteric effect has been unknown [33,34]. Our results indicate that weakened correlations of the nucleotide-binding regions in E241A^Gαt^ as a consequence of allosteric mutations in SIII/α3 and SII likely underlie the reported impaired GTP binding. Moreover, we identified residue E232^Gαt^ as a Gαt-specific primary contributor to the inter-lobe couplings in SIII, which has no direct counterparts in Ras or EF-Tu due to the absence of SIII (purple residues in **Table S4**). The simulation of mutation E232A^Gαt^ shows diminished couplings between PL, SI and SII, as well (**Figure S2A**). Similar effects of mutations R201A^Gαt^ and D234A^Gαt^ are also observed (**Figure S2B & C**).

Mutations of the counterpart residues E62A^Ras^ and K88A^Ras^ result in no significant change in the coupling of nucleotide binding loops in Ras (**Figure S1A**). Collectively these findings indicate that in Gαt and EF-Tu both N- and C-terminal α3 positions dynamically couple with SII, whereas in Ras the communication between α3 and SII is mainly through the C-terminus of α3. In addition, our results suggest that SIII plays a unique role in Gαt not only mediating the couplings between the two lobes but also allosterically maintaining the tight correlations between SI, SII and PL.

## Discussion

In this work, our updated PCA of Ras structures captures two new conformational clusters representing the GEF-bound state and “state 1”, respectively, in addition to the canonical GTP and GDP forms. By comparing the Ras PCA to PCA of Gαt/i and EF-Tu, we reveal common nucleotide dependent collective deformations of SI and SII across G protein families. Our extensive MD simulations and network analyses reveal common nucleotide-associated conformational dynamics in Ras, Gαt and EF-Tu. Specifically, these three systems have stronger intra-lobe1 (PL – SI and PL – SII) and inter-lobe (SII – SIII/α3) couplings in the GTP-bound state. Meanwhile, with the network comparison approach we further identify residue-wise determinants of commonalities and specificities across families. Residues M72^Ras^ (SII), V103^Ras^ (α3), D47/E49^Ras^ (L3) and R164^Ras^ (α5) are predicted to be crucial for inter-lobe communications in Ras. Mutations of these distal residues display decreased coupling strength in SI – PL. Interestingly, the analogous residues in the other two proteins, F211^Gαt^/I93^EF-Tu^ (SII), F255^Gαt^/V126^EF-Tu^ (α3), K188^Gαt^/R75^EF-Tu^ (L3) and D337^Gαt^/D207^EF-Tu^ (α5) also have important inter-lobe couplings and show similar decoupling effects upon alanine mutations. Besides the key residues that are common in the three systems, residues mediating inter-lobe couplings only in Gαt and EF-Tu are identified. These include R201^Gαt^/A86^EF-Tu^ and E241^Gαt^/Q115^EF-Tu^, whose cognates in Ras do not have significant effect on the nucleotide-binding regions upon mutation. In addition, Gαt specific residue E232^Gαt^ in SIII (which is missing in Ras and EF-Tu) is identified to be important to the couplings of the nucleotide-binding regions. Importantly, some of our highlighted mutants (D47A/E49A^Ras^, K188A^Gαt^, D207A^Gαt^ and R241A^Gαt^) have been reported to have functional effects by *in vitro* experiments. Our analysis provides insights into the atomistic mechanisms of these altered protein functions.

Using differential contact map analysis of crystallographic structures, Babu and colleagues recently suggested a universal activation mechanism of Gα [27]. In their model, structural contacts between α1 and α5 act as a ‘hub’ mediating the communications between α5 and the nucleotide. These contacts are broken upon the binding of receptor at α5, leading to a more flexible α1 and the destabilization of nucleotide binding. According to their studies, however, these critical α1/α5 contacts do not exist in Ras structures. Thus, they concluded that, unlike Gα, α5 in Ras does not have allosteric regulation of the nucleotide. It is worth noting that Babu’s work is purely based on the comparison of structures without considering protein dynamics. In fact, our study indicates that functionally important communications may not be directly observed from static structures. For example, the inter-lobe couplings between SII and SIII/α3 are not captured by PCA of structure ensemble, but they are clearly shown in our network analysis of structural dynamics. By inspecting structural dynamics, we find that α5 in Ras actually plays an allosteric role, in which point mutation (R164A) substantially disrupts the couplings in the nucleotide binding regions.

A previous study of Ras GTPases via an elastic network model – normal mode analysis (ENM-NMA) revealed similar bilobal substructures and found that functionally conserved modes are localized in the catalytic lobe1, whereas family-specific deformations are mainly found in the allosteric lobe2 [35]. The subsequent study via MD, in constrast, indicated that the conformational dynamics of Ras and Gαt are distinct, especially in the GDP state [36]. We note that in that study only a single MD simulation trajectory was analyzed, which is insufficient to assess the significance of the observed difference. Moreover, few atomistic details were given in that work. In our study, we make improvements by building ensemble-averaged networks based on multiple MD simulations instead of a single trajectory. This increases the robustness of the networks and largely reduces statistical errors. In addition, our correlation analysis provides residue wise predictions of potential important positions that mediate communications between functional regions. Overall, separation of functionally conserved and specific residues in conformational dynamics provides us unprecedented insights into protein evolution and engineering.

## Materials and Methods

### Crystallographic structures preparation

Atomic coordinates for all available Ras, Gαt/i and EF-Tu crystal structures were obtained from the RCSB Protein Data Bank [37] via sequence search utilities in the Bio3D package version 2.2 [38,39]. Structures with missing residues in the switch regions were not considered in this study, resulting in a total of 143 chains extracted from 121 unique structures for Ras, 53 chains from 36 unique structures for Gαt/i and 34 chains from 23 unique structures for EF-Tu (detailed in **Tables S1-S3**). Prior to analyzing the variability of the conformational ensemble, all structures were superposed iteratively to identify the most structurally invariable region. This procedure excludes residues with the largest positional differences (measured as an ellipsoid of variance determined from the Cartesian coordinate for equivalent Cα atoms) before each round of superposition, until only invariant “core” residues remained [40]. The identified “core” residues were used as the reference frame for the superposition of both crystal structures and subsequent MD trajectories.

### Principal component analysis

PCA was employed to characterize inter-conformer relationships of both Ras and Gαt/i. PCA is based on the diagonalization of the variance-covariance matrix, Σ, with element Σ_ij_ built from the Cartesian coordinates of Cα atoms, r, of the superposed structures:

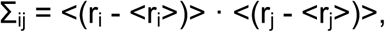

where i and j enumerate all 3N Cartesian coordinates (N is the number of atoms being considered), and <·> denotes the average value. The eigenvectors, or principal components, of Σ correspond to a linear basis set of the distribution of structures, whereas each eigenvalue describes the variance of the distribution along the corresponding eigenvector. Projection of the conformational ensemble onto the subspace defined by the top two largest PCs provides a low-dimensional display of structures, highlighting the major differences between conformers.

### Molecular dynamics simulations

Similar MD simulation protocols as those used in [18] were employed. Briefly, the AMBER12 [41] and corresponding force field ff99SB [42] were exploited in all simulations. Additional parameters for guanine nucleotides were taken from Meagher *et al.* [43]. The Mg^2+^·GDP-bound Ras crystal structure (PDB ID: 4Q21), Gαt structure (PDB ID: 1TAG) and EF-Tu structure (PDB ID: 1TUI) were used as the starting point for GDP-bound simulations. The Mg^2+^·GNP (PDB ID: 5P21), the Mg^2+^·GSP (PDB ID: 1TND) and the Mg^2+^·GNP (PDB ID: 1TTT) bound structures were used as the starting point for GTP-bound simulations of Ras, Gαt and EF-Tu, respectively. These structures were identified as cluster representatives from PCA of the crystallographic structures. Prior to MD simulations, the sulfur (S1γ)/nitrogen (N3β) atom in the GTP-analogue was replaced with the corresponding oxygen (O1γ) / oxygen (O3β) of GTP. All Asp and Glu were deprotonated whereas Arg and Lys were protonated. The protonation state of each His was determined by its local environment via the PROPKA method [ref]. Each protein system was solvated in a cubic pre-equilibrated TIP3P water box, where the distance was at least 12Å from the surface of the protein to any side of the box. Then sodium ions (Na^+^) were added to neutralize the system. Each MD simulation started with a four-stage energy minimization, and each stage employed 500 steps of steepest descent followed by 1500 steps of conjugate gradient. First, the atomic positions of ligands and protein were fixed and only solvent was relaxed. Second, ligands and protein side chains were relaxed with fixed protein backbone. Third, the full atoms of ligands and protein were relaxed with fixed solvent. Fourth, all atoms were free to relax with no constraint. Subsequent to energy minimization, 1ps of MD simulation was performed to increase the temperature of the system from 0K to 300K. Then 1ns of simulations at constant temperature (T=300K) and pressure (P=1bar) was further performed to equilibrate the system. Finally, 80ns of production MD was performed under the same condition as the equilibration. For long-range electrostatic interactions, particle mesh Ewald summation method was used, while for short-range non-bonded Van der Waals’ interactions, an 8Å cutoff was used. In addition, a 2-fs time step was use. The center-of-mass motion was removed every 1000 steps and the non-bonded neighbor list was updated every 25 steps.

### Correlation network construction

Consensus correlation networks were built from MD simulations to depict dynamic couplings among functional protein segments. A weighted network graph was constructed where each node represents an individual residue and the weight of edge between nodes, i and j, represents their Pearson’s inner product cross-correlation value cij [44] during MD trajectories. The approach is similar to the dynamical network analysis method introduced by Luthey-Schulten and colleagues [45]. However, instead of using a 4.5Å contact map of non-neighboring residues to define network edges, which were further weighted by a single correlation matrix, we constructed consensus networks based on five replicate simulations in the same way as described before [18].

### Network community

Hierarchical clustering was employed to identify residue groups, or communities, that are highly coupled to each other but loosely coupled to other residue groups. We used a betweenness clustering algorithm similar to that introduced by Girvan and Newman [46]. However, instead of partitioning according to the maximum modularity score, which is usually used in unweighted networks, we selected the partition closest to the maximum score but with the smallest number of communities (i.e. the earliest high scoring partition). This approach avoided the common cases that many small communities were generated with equally high partition scores. The resulting networks under different nucleotide-bound states showed largely consistent community partition in Ras, Gαt and EF-Tu, with differences mainly localized at the nucleotide binding PL, SI, SII and α1 regions. To facilitate comparison between states and families, the boundary of these regions was re-defined based on known conserved functional motifs. Re-analysis of the original residue cross-correlation matrices with the definition of communities was then performed. Only inter-community correlations were of interest, which were calculated as the sum of all underlying residue correlation values between two given communities satisfying that the smallest atom-atom distance between corresponding residue pairs was less than 4.5Å (for Gαt and EF-Tu) or 6 Å (for Ras) for more than 75% of total simulation frames. A larger cutoff was selected for Ras because the overall residue level correlations are weaker in Ras. A standard nonparametric Wilocox test was performed to evaluate the significance of the differences of inter-community correlations between distinct states.

## Acknowledgments

We thank Drs. John Tesmer and Guido Scarabelli for valuable discussions.

## Author Contributions

Conceived and designed the experiments: BJG. Performed the experiments: HL XQY, BJG. Analyzed the data: HL XQY, BJG. Wrote the paper: HL XQY BJG.

## Supporting Information

**Figure S1. Mutations of distal Gαt and EF-Tu specific residues perturb structural dynamics at nucleotide binding regions.**

In each panel, networks of wild type GTP-bound (WT-GTP, top) and mutant GTP-bound (MU-GTP, bottom) are compared. Red and blue edges indicate enhanced WT or MU couplings that are significantly (*p*-value <0.05). All other lines are colored gray. Specific mutations E241A^Gαt^ (**B**) and Q115A^EF-Tu^ (**C**) in α3 dramatically reduce the couplings between the functional regions PL, SI and SII, whereas the counterpart mutation K88A^Ras^ (**A**) has minor effects.

**Figure S2. Mutations of distal Gαt specific residues perturb structural dynamics at nucleotide binding regions.**

In each panel, networks of wild type GTP-bound (WT-GTP, top) and mutant GTP-bound (MU-GTP, bottom) are compared. Red and blue edges indicate enhanced WT or MU couplings that are significantly (*p*-value <0.05). All other lines are colored gray. Gαt specific mutations E232A^Gαt^ (**A**) in SIII dramatically reduce the couplings between the functional regions PL, SI and SII. Similar effects of mutations R201A^Gαt^ (**B**) and D234A^Gαt^ (**C**) are also observed in Gαt.

**Table S1. Analyzed crystallographic structures of Ras**

**Table S2. Analyzed crystallographic structures of G protein α subunit**

**Table S3. Analyzed crystallographic structures of EF-Tu**

**Table S4. Residue-wise contributions to inter-community couplings.**

The numbers represent the residue-wise contributions to inter-community couplings. For example, the sum of correlations between residue M72 in SII and all residues in SIII/ α3 is 1.19 (after filtering by contact map). The first row contains common counterpart residues (red) connecting SII and SIII/α3 in three proteins. The second row contains family-specific functional residues: residues in Gαt and EF-Tu (blue) contribute to the dynamic correlations between SII and SIII/α3, whereas their counterparts in Ras (green) have no contributions. The third row contains Gαt specific residue in SIII, which has no counterparts in the other two proteins.

